# Community density patterns estimated by species distribution modeling: the case study of an insect virus interaction

**DOI:** 10.1101/2024.02.08.579470

**Authors:** Stéphane Dupas, Jean-Louis Zeddam, Katherine Orbe, Barrera Cubillos Gloria Patricia, Laura Fernanda Villamizar, Patricia Mora, Jovanni Suquillo, Olivier Dangles, Aristóbulo Lopez-Avilla, Alba-Marina Cotes-Prado, Jean-Francois Silvain

## Abstract

1. Time delays complicates the analysis of trophic dependence, which requires large time series data to study local associations.
2. Here we propose using species distribution modeling. This approach removes confounding time lag effects and allows using data obtained separately in the different species.
3. Since the approach is correlative, it cannot be interpreted in terms of causality.
4. We apply the method to the interaction between the invasive potato moth Tecia solanivora and its granulovirus PhoGV in the Northern Andes. Host density was analyzed based on 1206 pheromone trap data from 106 sampled sites in Ecuador, Colombia and Venezuela. Virus prevalence was evaluated in 15 localities from 3 regions in Ecuador and Colombia. glm models were optimized for both variables on bioclimatic variables. Predicted virus prevalence was not significantly correlated to host density in the sampled virus sites. Across the climatic range covered by the study, correlation was R=−0.053. Of the total population of insect in this range, 26% were expected to be infected.
5. Infection status was also analyzed for spatial structure at different scales: storage bag, storage room, field, locality, country. Locality and storage bag explained respectively 8% and 26% of the total deviance in infection status in glm analysis. Field and storage structure differed within locality but not always in the same direction.
6. This basic method may help studying statistical relationships between species density across a number of trophic models making use of existing non sympatric data, with none or limited additional sampling effort.

## Introduction

The characterization of species interactions is critical to predict ecosystem dynamics and stability [1]. An array of techniques have been developed to infer them from the analysis of time series [2–7]. Time lags in species response is among the confounding factors. Theory predicts 1/4-period lags in consumer resource abundance, and many datasets even show consumer-resource oscillations with an antiphase (1/2-period lags) or nearly antiphase period due to eco-evolutionary dynamics [8,9]. These time lags can yield positive or negative correlations in contemporaneous samples. Inference of both time lag and interaction strength evolution relies long term time series data [10,11], analysed based on local similarity analysis [7], empirical dynamic modeling [5], and extensions to track non linear responses [3] and changes [4]. Causality is still not inferred due to latent confounders but statistical issues can be handled. In some highly evident situations, like insect pathogens epizootics, it is possible to relate pathogen to host density. But for quantitative tests, apart from systems of major importance for their ecosystem services for which important research can be funded, it is relatively difficult to get sufficient data to retrieve information on food web structure and interaction strength variations.

Strydom et al. [6] proposed a neural network model to estimate average interactions across space and time. This analysis cannot replace more mechanistic time series analyses described above but may provide an important parameter for decision support for species management : in many management situations, causality is not the only issue, observing a correlation may still have management relevance within the study area. In this study, we propose a very simple approach to estimate spatial correlations in the time average of species abundance by using niche models estimates. Niche modeling predicts average densities of interacting species based on environmental variables. Adding environmental driver data should improve the power compared to inference based on species data only such as Strydom’s neural networks.

Insect viruses are good example of simple system models where data on environmental drivers of communities is lacking. They are expected to have important ecological impact on their instect host populations in the wild [12]. This potential impact stimulated their use for controlling insect pests in agriculture [13,14]. They are usually used with an inundative rather than inoculative strategy [15]. But resident populations of insect virus may have naturally tremendous impact on insect populations, considering their prevalence and diversity [16,17]. However, ecological data on their natural interactions and impact as natural agents is still scarce. There is little estimation of top down or bottom up forces or even correlation patterns in these systems. As for fungus [18], a better knowledge of their epidemiology and distribution should help improving their use in biological control of insect pests, and should ensure a higher sustainability of their deleterious effects on host pest populations.

We developed a methodological framework for testing patterns of pathogen prevalence on host density in *Tecia solanivora* (Lepidoptera: gelechiidae) (Povolny) potato tuber moth host, and its granulovirus (Baculoviridae) interaction. *T. solanivora* is an invasive species from Guatemala that extended its distribution area to Northern Andean region down to Ecuador [19–21]. As this species is invasive, interaction with most natural or introduced biological control agents is recent in Northern Andes. The biological control of potato tuber moth (PTM) using insect virus has a longstanding history [22]. *Phthorimaea operculella* granulovirus or PhoGv (Genus: Alphabaculovirus; Family Baculoviridae) was initially discovered in Australia [23]. It is now used worldwide for the biological control of PTMs. Many studies have been published dealing with the use of this virus in pest control [22,24], pathogenicity[24–27], genetic variations [28–30], histopathology [31], persistence in the soil [32], co-infection [33], among others. However, little information is available on the epizootiology of this virus [34–36].

The aim of this work was first to determine the level of infestation of *Tecia solanivora*, by *PhopGV* in the field and then to analyze the virus prevalence in relation to the density of the insect pest across altitudes and between field and storage facilities in Northern Andes (Ecuador and Colombia). Furthermore, we propose an original approach using niche models to average temporal variations of host for the estimation of correlation patterns for integrated pest management.

## Materials and methods

### Insect data

*T. solanivora* densities were evaluated using pheromone baited traps located along altitudinal gradients of larger amplitude than for virus prevalence in Ecuador, Colombia and Venezuela. Five different surveys, representing different regions and/or periods of time, were carried out (Table 1). A total of 1216 pheromone trap samples from 106 localities were obtained across Northern Andes (Fig 1).

**Table 1.** Description of pheromone trap surveys used in this study.

| Region | Nu<br>mber of<br>localities | Peri<br>od | Fre<br>quency | Nu<br>mber of<br>samples | Ref |
| --- | --- | --- | --- | --- | --- |
| Central<br>Ecuador | 42 | Jan<br>2006 -<br>Dec<br>2006 | 3<br>weeks | 714 | Dan<br>gles et al,<br>2008 |
| Southern<br>Ecuador | 1 | Apr<br>2008 | 3<br>days | 1 | This<br>study |
| Central<br>Ecuador | 14 | Feb<br>2008 – Apr<br>2008 | 3<br>weeks | 9 | This<br>study |
| Northern<br>ecuador | 7 | Sep<br>2009 – Oct<br>2009 | 1<br>week | 54 | This<br>study |
| Colombia | 18 | July<br>2008; Sep-Dec 2009 | 4<br>days | 96 | This<br>study |
| Venezuel<br>a | 25 | June<br>1984 – Nov<br>1984 | 1<br>week | 330 | Sala<br>zar & Torres<br>2006 |
| Total | 107 |  |  | 1204 |  |

**Fig 1:**
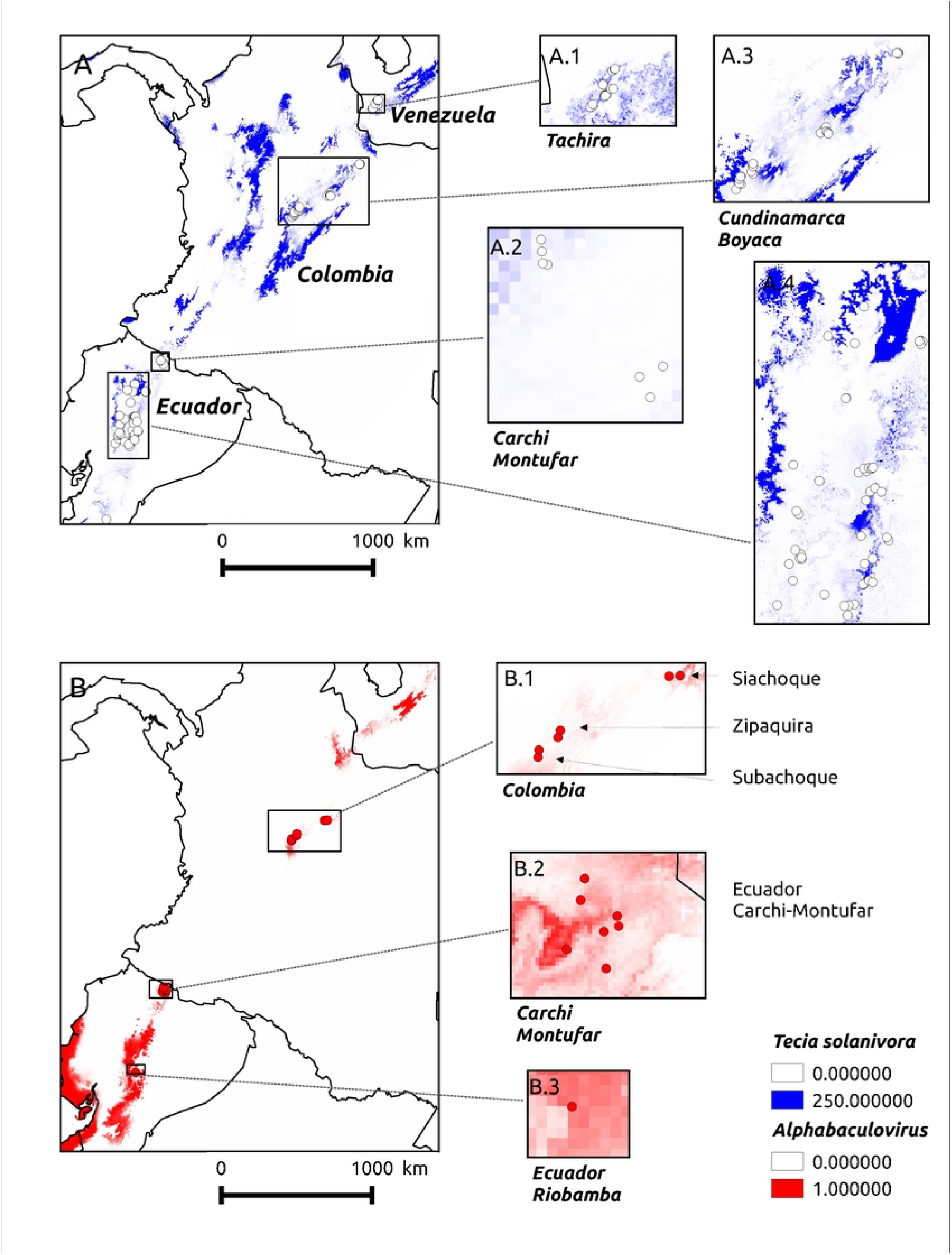
Sampling sites and predicted distributions of *PhopGV* and *Tecia solanivora*. PhopGV prevalence estimated by PCR.

### Virus occurrence data

*T. solanivora* larvae (517) were collected from infested tubers from 15 localities dispatched along 5 altitudinal gradients, three located in Colombia from 2800 to 2940 masl Subachoque, Siachoque and Zipaquira and three in Ecuador from (Montufar-Carchi and Riobamba) (distribution of localities in presented in figure 1). In these localities, the granulovirus had never been applied for biological control purposes. Tubers were collected and dissected in the laboratory. Healthy larvae and/or larvae with signs of granulovirus disease were individually collected and conserved in alcohol for DNA analysis. Larval tissues were homogenized in 50 μL bidistilled H2O and centrifuged at 1000rpm for 5 min at 4ºC. To recover OBs, supernatants were centrifuged at 15000 rpm for 15 min and pellets resuspended in 100 μL bidistilled H2O. Suspensions were incubated with 100 μl vol. of 0,5M Na2CO3, 50 μL SDS 10% for 10 min at 60ºC and centrifuged at 5000 rpm for 5 min. The supernatants were digested with 25 μL proteinase K (20mg/mL) at 56ºC for one hour and DNA was extracted using a phenol/chloroform/isoamyl alcohol protocol and then precipitated with ethanol, as described in previous works [37].

DNA was extracted using proteinase K digestion, The presence of the *PhopGV* was detected by PCR using primer pair, 83.2 5’-CCGCGCCGATTACCAACAGCAGC-3’ and 84.1 5’-GAACTGTTAAACGGCTTGAGTGAGCG-3’ designed over the GenBank sequence JX170206.1.

on 517 *T. solanivora* collected larvae, amplifying a 241 bp region encompassing part of gene 83 and 84 of the PhoGV genome. Amplification conditions were: 94ºC for 1min, 30 cycles (94ºC for 1min, 50ºC for 1min, 72ºC for 1 min), and a final extension of 7 min. Conditions were optimized in Colombia and Ecuador separately to reduce false negatives. Colombian samples were amplified in Corpoica-Tibaitata, Colombia using the following 10X mix: dNTPs 200 μM (Pharmacia Biotech 27-1850), 0.5 μM of both primers, MgCl2 2.0 mM, buffer 10X (50 mM KCl, 10 mM Tris HCl pH 9.0 0.1% Nonidet), 2U Taq polymerase (Promega M1665). Ecuadorian samples were amplified in Santa Catalina: PCR mix contained dNTPs 250 μM (Invitrogen, reference?), 0.5 μM of both primers, MgCl2 1.6 mM, buffer 10X (50 mM KCl, 20 mM Tris Hcl, pH 8.4), 0.075U/ul Platinium Taq polymerase (Invitrogen).

All data files are available from the mendeley database (accession number p99dbv75c4/1).

### Species distribution models

Generalized linear models (*glm*) were optimized for *T. solanivora* abundances assessed by pheromone baited traps, and for PhoGV occurrence assessed by PCR test. Probability distributions were negative binomial with *log* link function for *T. solanivora* data and binomial with *logit* link function for PhoGV data. Independent variables were the region of collection (Central Ecuador, Northern Ecuador, Central Colombia, Venezuela), altitude and bioclimatic variables {Citation}].

Bioclimatic variables were obtained by overlaying the 30” (about 1 km) bioclimatic layers onto sites coordinates. Sites differing by more than 200m between WorldClim and measured altitude were removed from the analysis[39]. For the insect density, square root of altitude and all bioclimatic variables were included. For virus prevalence, only BIO1 (mean temperature), BIO6 (minimum temperature of the coldest month) and BIO12 (mean precipitation) were included. We built generalized linear models using a full backward stepwise approach using step function of R with BIC criterion for variable selection [40]. We assumed negative binomial distribution of the response and log link function for the insect and binomial distribution of the response and logit link function for the virus. Distribution maps were reconstituted from the selected model predictions using WorldClim bioclimatic layers in a 30 arc second resolution (< 1 km). The relationships between insect and virus distributions were analysed. The cell grid of the map used for the analysis were those having all bioclimatic and altitude values within the range of the sites sampled for the virus, and for the insects. Parasite prevalence and pest density were estimated by the model in these sites. The correlation between virus prevalence and host density were plotted and calculated using *glm* model analysis of deviance. For significance testing, only localities where virus had been sampled were considered. Overall prevalence was calculated as the total number of infected insects divided by the total number of insects in the range considered, using equation

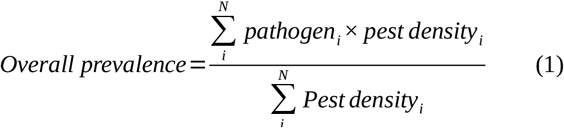

Where *i* and *N* are the grid cell number and number of grids cell satisfying climatic conditions of the sampled area, respectively. The data were analyzed in R 3.2.2. The script is available as S1 File. R libraries *raster, sp, mapplots* and *rgdal* were called in the script.

### Hierarchical structure of virus prevalence

Autocorrelation of virus prevalence at different scales (country, locality, site, storage room/field, potato bag and tuber) were analyzed with *glm*. Binomial distribution and *logit* link function were assumed. First model tested country and locality effects (locality nested in country). Then, nested *glm* tested the effect of locality, storage bag and tuber (tuber nested in bag nested in locality). The third model tested differences between field and storage room within locality.

## Results

### Hierarchical structure

*PhopGV* prevalence was estimated on 517 larvae from 15 sites in Colombia and Ecuador using PCR detection. Important variations were observed among sites (Figure 1). Prevalence was higher in Ecuador (29%) than in Colombia (4%) (Figure 1, Figure 2). In Ecuador, virus infection variation was analyzed in relation to locality, field versus potato storage rooms, potato storage bag, and potato tuber (Table 1). There were significant effects of Site (p<0,0001, 53.6% of the deviance, df=5) and potato bag (p<0.05, 17.2% of the deviance, df=12). Differences between potatoes within bag were not significant. Difference between storage room and field were not significant. In the two sites where both storage rooms and fields have been sampled, the difference was not in the same direction: there was significantly more viral infections in the field in one locality (Chutan bajo, Montufar, Carchi, logit +4.6, p<0.01) and significantly less infections in the field in the other locality (El Chamizo, Montufar, Carchi, logit −3.5, p<0.05).

**Figure 2:**
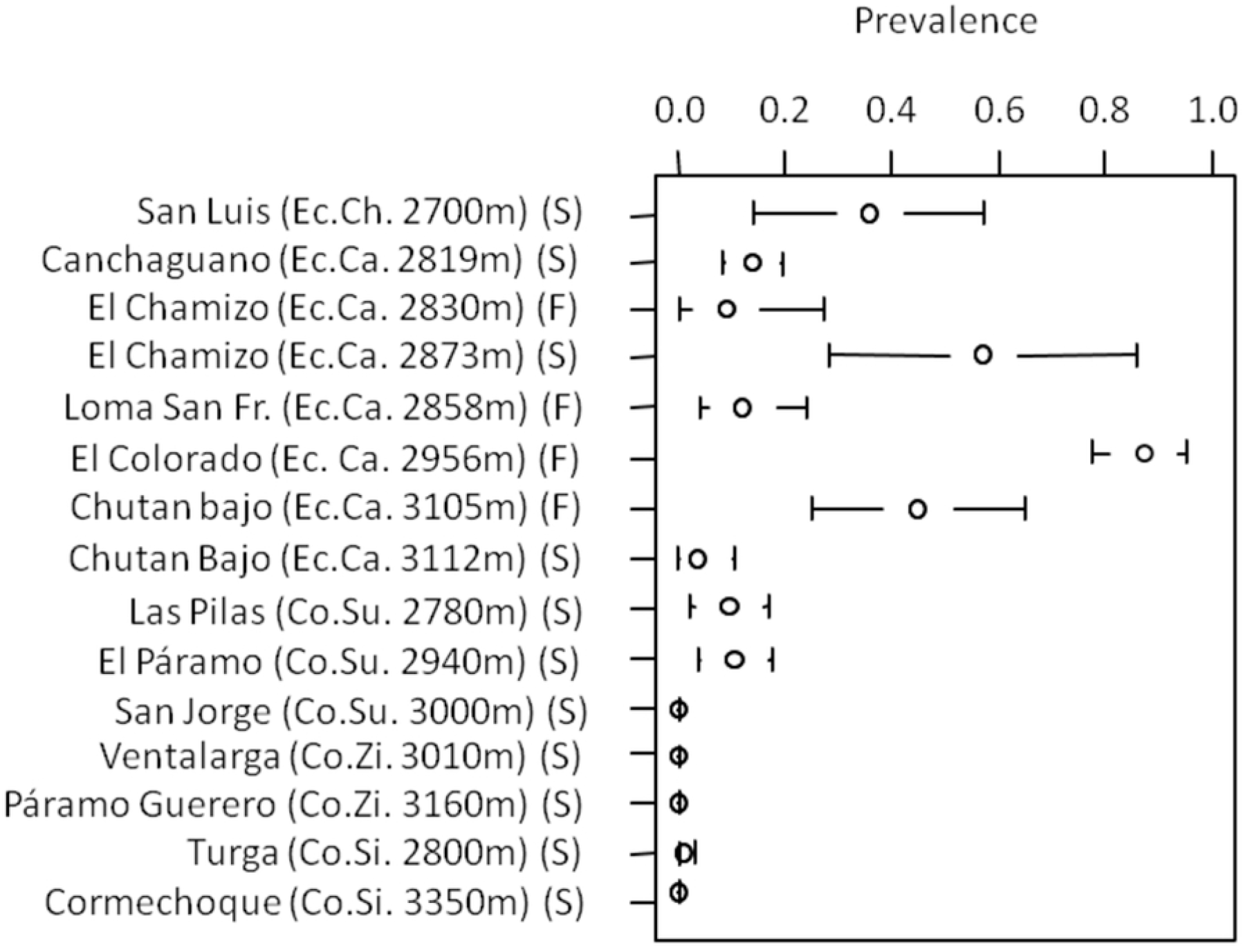
PhopGV prevalence estimated by PCR. Country, gradient and altitude are given between brackets. Co. : Colombia. Ec.: Ecuador. Ch. Chimborazo: Ca. Carchi: Su.: Suachoque, Zi.: Zipaquira, Si.: Siachoque. S: Storage room sampling. F: Field sampling.

### Average correlations

Species distribution models were optimized for the two partners using field occurrence data for the virus (517 moth across 13 localities in Ecuador and Colombia) and field abundance data for the insect (1203 pheromone trap data across 105 localities in Ecuador, Colombia and Venezuela). Stepwise *glm* results are presented in S1 Table and S2 Table for moth and virus, respectively.

In each locality sampled for the analysis of *PhopGV* larval prevalence, the density of *T. solanivora* was estimated based on the *glm* regression predictor. Altitude, temperature at the driest month and average precipitation all had significant negative effect on prevalence (p<0.0001), precipitation being the most significant (p<10^−16^) (S2 Table). Projected moth and virus abundance are presented in figure 1. The relationship between log potato moth species densities predicted by niche modeling and virus prevalence predicted by niche model within altitudinal gradients are represented in figure 2 for the sites where virus were sampled. The prevalence of the virus was not significantly dependent on host density (R=0.182 NS, figure 2). To have a broader view (without statistical test), we studied the relationship between all the cells of the map within the ranges of the virus localities and within the ranges of the *T. solanivora* survey localities for all the variables used in their respective models (S1 Fig. for the relationship, and S1 Fig. for the corresponding cells in the map). Correlation was close to zero in these cells (R=−0.053). The overall percentage of insect infected by the virus in this climatic range was 23.6%.

## Discussion

In this study, prevalence of the baculovirus *PhopGV* was assessed in populations of three potato moth species found in farmers’ potato storage facilities located at different altitudes in the Northern Andes. While no symptomatic *PhopGV*-infected PTM larvae was found during the survey, the virus was however relatively frequent among the sampled individuals from Colombia and Ecuador but as covert infections (23.6% prevalence overall). These observations on virus prevalence in *T. solanivora* are in accordance with previous reports limited to *PhopGV* in *P. operculella* larvae in Australia and Tunisia [34,35] and with prevalences observed up to 50% in granulovirus in general [41]. In the case of *PhopGV*, it seems that sublethal infections are the rule while symptomatic larvae are not frequently found. Similarly, epizootics, if any, would be exceptional.

For each altitudinal gradient, no significant relationship was observed between *PhopGV* prevalence and density of the insect *T. solanivora* (Figure 2). The absence of significant relationship between host density and virus prevalence suggests that environmental drivers of moth and virus are independent [42]. For the virus, our result suggest a significant negative effect of altitude, temperature or the driest quarter and precipitation. These results are consistent with [43] in its recent review of soil baculovirus reservoir. Negative effect of UV (related to altitude and temperature in the Andes) on virus transmission was presented as commonly accepted and was again recently reported to reduce latent baculovirus transmission across crop seasons[44]. Moist soil, consistent with our precipitation effect, were also reported to reduce baculovirus activity. Finally elevated temperatures >60°C do inactivate occlusion bodies within few minutes [43]. With more data, more factors could be include in our model, and the direct effect of climatic factors on PhoGV transmission could be distinguished and separeted from host density effect.

We observed autocorrelations of virus prevalence at different scales (tubers, bag, storage room/field, site, and region). The autocorrelations observed suggests that bag storage contribute to *PhopGV* epizooty. Transmission of the virus may be increased by keeping the same bags for storage from one season to the other, and also, possibly by leaving storage residuals within the room between seasons. Recycling of used bags after the tubers are sold to the market may also favor virus transmission at larger distance. As a conclusion, the same factor which favor the dispersal of PTM at large distance may favor the dispersal of *PhopGV* [22,45–47].

Measuring food web patterns and processes is a major focus of ecology [42,48,49]. Our methodology is relatively simple and allows to measure average density relationships between trophic levels across geographic range of species. It does not require occurrence data to be obtained at the same time and place for each species of the food web, facilitating meta-analysis approach. It should therefore be possible to use much larger data bases than in the current meta-analysis which are restricted to data where multiple trophic levels are studied together across large spatial and time scales [48]. Niche model density estimates have however to be considered with caution and not used outside the calibration range of the model where they may not be valid [50]. In our work, we therefore removed sites outside the data climatic and geographic range to estimate correlations.

Another issue is significance testing. A conservative test would require the number of sites used for correlation testing not to exceed the number of sites of any of the species analyzed in the test. In our work, we therefore used the sites of the species having the least number of sites (the virus) for correlation testing. In the context of integrated pest management strategies, habitat management variables could be added to the environmental dataset to study jointly their effects on pest and beneficial insects or control agents. The method is in theory applicable to any consumer resource or food web system, and in practice should be valuable to assess a number of agents for biological control. The estimation of average density at each trophic level depending on nutrient resource or climate may provide data for a more general view of the consequences of global changes in climate and energy and matter flows on food web structure.

## Aknowledgements

This work has been founded by ECOFOR program ENTOAND 2007-2011.

## Supporting information

**S1 Fig. *Tecia solanivora* predicted density and PhoGV predicted prevalence, by the glm models**. Presented are the cells of the map where predictor variables fall in the range where both Tecia solanivora and PhoGV have been sampled.

**S1 Table. Stepwise polynomial generalized linear model regression procedure using BIC criteria for potato tuber moth in the north Andean region**. (A) *Phthorimmaea operculella*, (B) *Tecia solanivora*, and (C) *Symetrischemma tangolias*. Full model comprised following dependent variables : Alt = Alitude, BIOi = worldclim bioclimatic variables (see details below), their squared values I(.)^2, and the belonging to different regions: Central Ecuador (Chimborazo, Bolivar, Tungurahua, Cotopaxi, and Pichincha provinces), Northern Ecuador (Carchi), Central Colombia (Cundinamarca, Boyaca), and Venezuela. BIOi : BIO1 = Annual Mean Temperature, BIO2 = Mean Diurnal Range (Mean of monthly (max temp - min temp)). BIO3 = Isothermality (BIO2/BIO7) (* 100), BIO4 = Temperature Seasonality (standard deviation *100), BIO5 = Max Temperature of Warmest Month, BIO6 = Min Temperature of Coldest Month, BIO7 = Temperature Annual Range (BIO5-BIO6), BIO8 = Mean Temperature of Wettest Quarter, BIO9 = Mean Temperature of Driest Quarter, BIO10 = Mean Temperature of Warmest Quarter, BIO11 = Mean Temperature of Coldest Quarter, BIO12 = Annual Precipitation, BIO13 = Precipitation of Wettest Month, BIO14 = Precipitation of Driest Month, BIO15 = Precipitation Seasonality (Coefficient of Variation), BIO16 = Precipitation of Wettest Quarter, BIO17 = Precipitation of Driest Quarter, BIO18 = Precipitation of Warmest Quarter, BIO19 = Precipitation of Coldest.

**S2 Table. Stepwise polynomial generalized linear model *S2 Table. Stepwise polynomial generalized linear model regression for pohGV prevalence in Tecia solanivora. Alt = Alitude, BIO9 = Mean Temperature of Driest Quarter, BIO12 = Annual Precipitation***.. Alt = Alitude, BIO9 = Mean Temperature of Driest Quarter, BIO12 = Annual Precipitation.

***S2 File. R Script for the analysis of relationships between PhoGV and Potato tuber moth distributions***.

